# A Vertically and Horizontally Transmitted RNA Virus Facilitates Egg Hatching of a Parasitoid Wasp

**DOI:** 10.1101/2024.10.30.621033

**Authors:** Y. Izraeli, G. Wodowski, N. Mozes-Daube, J. Varaldi, E. Zchori-Fein, E. Chiel

## Abstract

Information on the impacts of RNA viruses inhabiting insect hosts is scarce. Here, we studied the effects of a recently described RNA virus, termed AnvRV, on its host, the parasitoid wasp *Anagyrus vladimiri* (Hymenoptera: Encyrtidae), an important natural enemy of mealybug pests. AnvRV was found to be maternally transmitted with very high fidelity but not paternally. Additionally, AnvRV was horizontally transferred at an efficiency of 23% from infected to uninfected wasp larvae that develop together inside the same mealybug host (superparasitism). To test the effects of AnvRV on *A. vladimiri*, the virus horizontal transmission was utilized to establish AnvRV-infected (RV^+^) and uninfected (RV^-^) isogenic wasp lines, a method rarely applied and novel to RNA virus-parasitoid systems. Longevity, developmental time, sex ratio, and fecundity of RV^+^ and RV^-^ *A. vladimiri* were very similar. Nonetheless, the egg hatching rate of RV^+^ wasps was markedly and significantly higher than that of RV^-^ wasps, especially in hosts that were not superparasitized. Additionally, less encapsulation marks (the main form of mealybug immunity) were found around RV^+^ eggs inside parasitized mealybug hosts. Taken together, the data suggest that AnvRV is affecting the mealybugs’ physiology in a way that improves first stages of wasps’ development. These findings present a rare example of interaction between an RNA virus and a parasitoid and may provide a tool for the improvement of biological control efforts.

## Introduction

Many endosymbiotic microorganisms form successful long-term relationships with insects, and have significant effects on various biological features, ranging from parasitism to obligate mutualism (Zchori-Fein and Bourtzis, 2011). While insect-bacteria and insect-fungi interactions have been extensively studied, viruses associated with insects, especially RNA viruses, have received less attention (Nouri et al., 2018), despite their ubiquitous association with insects (Shi et al., 2016; Gilbert and Belliardo, 2022). Most of the research on insect viruses has focused on pathogens, yet many insect species harbor a diverse range of non-pathogenic viruses (Bonning, 2020; Wu et al., 2020; Guinet et al., 2024; Varaldi et al., 2024). One of the best-studied examples of insect mutualistic viruses are polydnaviruses (PDVs), which are endogenized viruses found in some clades of parasitoid wasps within the Ichneumonidae and Braconidae families. PDVs are injected with the wasp eggs into their lepidopteran host where they suppress the host’s immunity, thereby promoting the successful development of the wasp offspring (Herniou et al., 2013; Strand and Burke, 2013). While PDV genomes have integrated into the wasp genome, parasitoids can also inject non-integrated DNA or RNA viruses together with their eggs into the host, thereby increasing their parasitism success. Examples include the DIEPV – a poxvirus in the fruit fly parasitoid *Diachasmimorpha longicaudata*, the ascovirus DpAV4 in the leek moth parasitoid *Diadromus pulchellus*, and the RNA virus DpRV2 in *D. pulchellus* (Renault et al., 2003; Coffman et al., 2022, 2024). Some viruses alter the parasitoids’ host behavior, for example in the solitary parasitoid wasp *Leptopilina boulardi*, LbFV induces superparasitism (*i*.*e*., laying eggs in already parasitized hosts) thus promoting the virus’s horizontal transmission within the superparasitized host (Varaldi et al., 2003, 2006). Additionally, viruses can manipulate the sex ratio of their hosts by inducing male killing, e.g. *Partitivirus* in *Drosophila* (Kageyama et al., 2023) and in the tea moth (Nakanishi et al., 2008; Fujita et al., 2021). Evidently then, viruses have diverse and substantial effects on the ecology of insects.

Here, we present a study on the effects of a recently discovered virus on its host, the parasitoid wasp *Anagyrus vladimiri* Triapitsyn, (Hymenoptera: Encyrtidae) (previously named *A. pseudococci*). *Anagyrus vladimiri* is an endoparasitoid wasp which is often used in biological control programs of two major global pest species: the citrus mealybug, *Planococcus citri* and the vine mealybug *P. ficus* (Hemiptera: Pseudococcidae) (Bugila et al., 2015). The adult female parasitoid lays a single egg into the mealybug’s body, the hatching wasp larva feeds and develops inside the host, pupates and eventually the adult emerges from the mummified host (Avidov et al., 1967). However, within the mealybugs’ hemocoel, the parasitoids’ egg may be encapsulated by immune hemocytes, which adhere to the surface of the egg, forming a multicellular melanized capsule-like envelope around it, usually eliminating it (Blumberg, 1997; Suma et al., 2012). As a counter adaptation, although only one wasp can complete its development in the host, *A. vladimiri* often lay eggs in already parasitized hosts, a behavior termed superparasitism, thereby increasing the chance of getting an offspring from the host (Islam and Copland, 2000; Suma et al., 2012). Both superparasitism behavior and the encapsulation response are quite common in parasitoids, and, as mentioned above, in some species these phenomena have been reported to be modulated by viruses (Herniou et al., 2013; Martinez et al., 2012; Varaldi et al., 2003, 2006).

Recently, we documented a new Reovirus in *A. vladimiri*, termed AnvRV. It is a double strand RNA virus belonging to the family *Spinareoviridae*, order *Reovirales* (previous family: *Reoviridae*) (Izraeli et al., 2022). The facts that AnvRV was not detected in unparasitized mealybugs, and is found in the ovaries of female wasps, suggest vertical transmission. Importantly, AnvRV-carrying *A. vladimiri* wasps in the lab do not exhibit any disease symptoms, raising questions regarding the phenotypic effects and mechanisms by which AnvRV is maintained in *A. vladimiri* populations. Here, these questions are addressed by: **a**) studying the transmission routes of AnvRV in *A. vladimiri;* **b**) establishing genetically-identical *A. vladimiri* lines with and without AnvRV, **c**) studying the phenotypic effects of AnvRV on *A. vladimiri*, including superparasitism behavior and encapsulation.

## Materials and Methods

### Origin and rearing of insect lines

All information on the *Anagyrus vladimiri* lines used in this study are detailed in Izraeli *et al*. 2022. Briefly, in 2019 a line of *A. vladimiri* was obtained from a mass-rearing facility, fed with antibiotics to remove bacterial symbionts (mainly *Wolbachia*). This line, termed Lab-RV^+^, stably harbors AnvRV. A second *A. vladimiri* line was established from wasps collected in a vineyard in northern Israel during the summer of 2020. *Wolbachia* and AnvRV are absent in this wasp line, which is hereafter referred to as ‘Field-RV^**-**^’ line (or simply ‘RV^-^’). Both lines were reared in the lab on the citrus mealybug *Planococcus citri*, which were fed on sprouted potatoes. The parasitoids maintenance and the experiments took place under controlled conditions of 26±1 °C, 60 ± 20% RH, and 16L:8D photoperiod regime.

### RT-PCR detections of AnvRV

RNA was extracted from individual wasps (RNeazy plus kit, Qiagen) and reverse transcribed to cDNA using the RT-PCRbio kit (PCR Biosystems). cDNA samples were used as templates for diagnostic PCRs with AnvRV-specific primers (K27_reo_F, forward: 5’-CAAACACGGCTCAAATGGCA-3’; K27_reo_R, reverse: 5’-TGAGTAGCGTCCTGATGGGA-3’). The PCR conditions were: 94 °C 30’’, 60 °C 30’’, 72 °C 60’’ (34 cycles), and final elongation at 72 °C for 5 min.

### Vertical transmission (VT) of AnvRV in Anagyrus vladimiri

Studying the effects of microbial symbionts on their hosts requires a comparison between host lines sharing identical, or at least very similar, genetic background, but varying in symbiont composition. Information on both maternal and paternal vertical transmission (VT) rates of AnvRV is crucial for the establishment of stable parasitoid lines that can serve for phenotype experiments.

To determine VT efficiency, ∼80 parasitoid pupae (inside their mummified mealybug hosts) from each of the Lab-RV^+^ and the Field-RV^-^ lines were isolated in Eppendorf tubes, one mummy per tube, to avoid un-controlled mating. The newly emerged virgin wasps were randomly assigned to the four possible crosses: RV^+^♀/ RV^+^♂, RV^+^♀/ RV^-^♂, RV^-^♀/ RV^+^♂, RV^-^♀/ RV^-^♂, a setup that allowed to test and quantify both maternal and paternal transmission. Each couple was placed in a 30ml plastic cup covered with fabric mesh, with ∼50 mealybug hosts. After 48h the wasps were removed from the cup and their infection status was verified by PCR, as described above. The mealybugs were incubated until they mummified, and 12 mummies from each cup were isolated to Eppendorf tubes to avoid possible horizontal transmission between siblings. Between 4-8 offspring (females and males) of each cup were tested by diagnostic PCR for the presence of AnvRV.

### Horizontal transmission (HT) of AnvRV and establishment of RV^+^ and RV^-^isogenic wasp lines

To test whether AnvRV can be horizontally transmitted (HT) between adult wasps (*i*.*e*., by mating or physical contact), and/or between larvae developing together in the same mealybug host (superparasitism), the arrhenotokous reproduction of *A. vladimiri* was utilized. The sex determination mechanism in all Hymenopteran species (bees, wasps and ants) is arrhenotoky: unfertilized eggs develop into haploid males and fertilized eggs develop into diploid females, therefore virgin females can produce only male (haploid) offspring. To maximize the chance that females will superparasitize the mealybugs, three **virgin** Lab-RV^+^ and two **mated** Field-RV^-^ (‘RV^-^’) females were placed together in cups with 25 hosts only for 48h. Hence, female offspring in this experiment are inevitably the daughters of the Field-RV^-^ mothers, as virgin Lab-RV^+^ were able to produce only males. After 48h, all adult females were removed and PCR tested to assess HT between adult wasps (if HT exists, we expect more than three positive females). The mealybug hosts were incubated until they mummified (∼ 9 days), and 12 mummies from each cup were isolated in Eppendorf tubes to avoid possible horizontal transmission between siblings and uncontrolled mating. Upon emergence, F_1_ females were singly backcrossed with males from the RV^-^ line, allowed to oviposit on new hosts for three days, and then tested by PCR. This setup was carried out twice (*n*=5 in the first experiment, *n*=10 in the second experiment). Offspring (F_2_) of replicates that were found to be AnvRV-positive were used to establish a new AnvRV^+^ line, termed hereafter ‘Field-RV^+^’ (or simply ‘RV^+^’) line, having identical genetic background as the Field-RV^-^ line (see Fig. 1 for setup illustration).

**Fig. 1.**
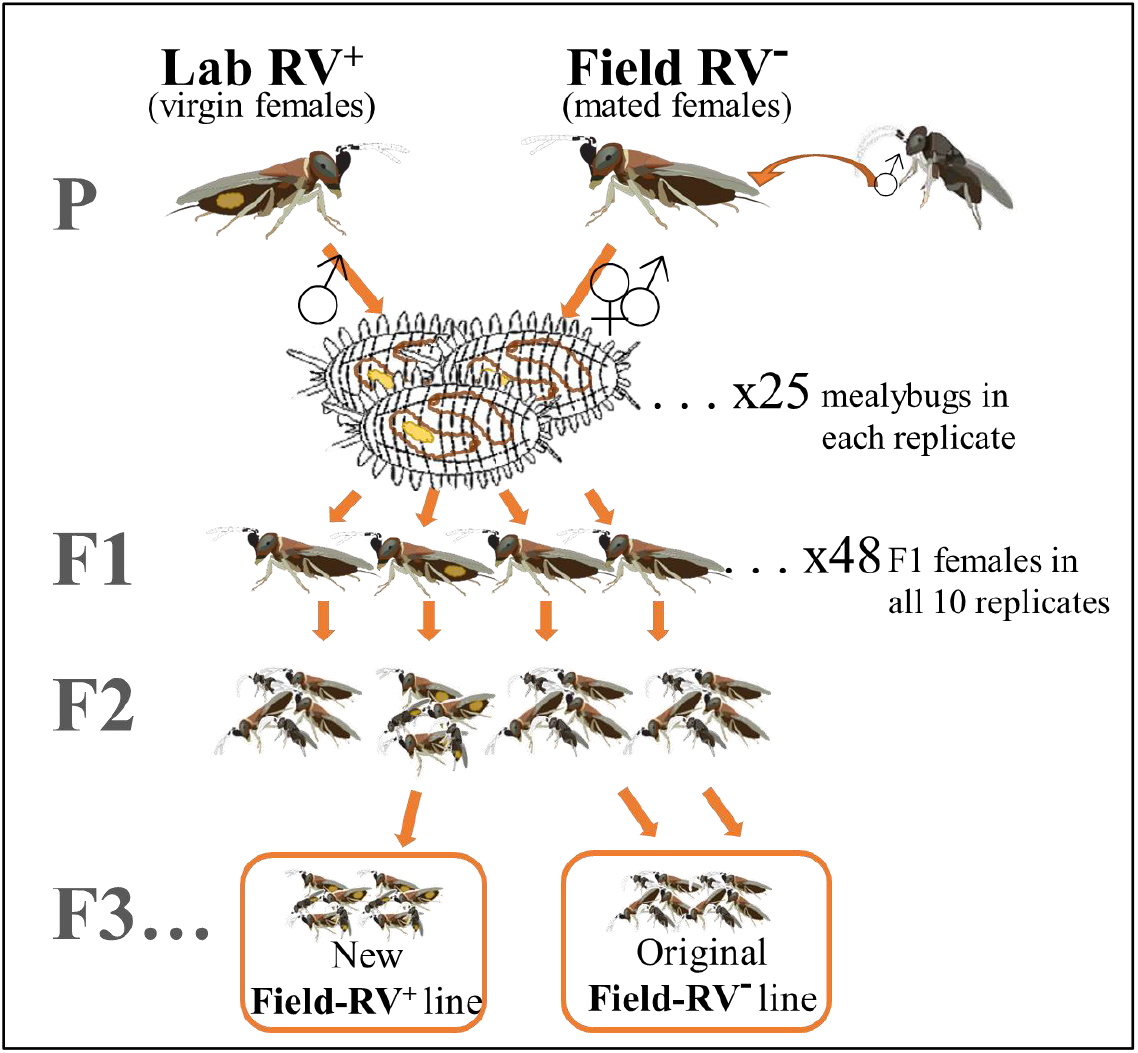
Setup of HT experiment and establishment of the new Field-RV^+^ wasp line. Yellow spots on wasps indicate that they are AnvRV-infected. Large letters on the left denote the generation, starting from P (parental) to F3… (ongoing reproduction of the established line). ♀ and ♂ symbols denote female and male progeny of the virgin and mated P wasps. These progeny larvae are competing with each other within the bodies of mealybug hosts. Insect icons courtesy of ‘BioBee Sde Eliyahu Ltd’. See Methods section for more details.

### Fitness experiments

To test whether AnvRV affects developmental time, longevity, fecundity, or sex ratio of *A. vladimiri*, these fitness parameters were measured and compared between the newly established RV^+^ line and the RV^-^ line.

Twenty pairs of 0-24 h old female and male wasps (P generation) of each line, were placed in a ventilated cup with a potato sprout infested with ∼20 mealybugs, one couple per cup. To obtain even-aged wasp eggs so that developmental time could be determined, the wasps were removed from those cups (marked ‘A’) after 6 h. The wasps were then transferred to new (‘B’) cups to continue ovipositing until they die. Every two days, fresh mealybugs were added *ad libitum* to the ‘B’ cups, to allow females to lay as many eggs as they can. Mortality of the females was recorded daily to determine longevity and survival rate. Emergence of F_1_ wasps in the ‘A’ cups was recorded twice daily to determine the developmental time. All F_1_ emerging wasps were counted to determine fecundity and sex ratio (a sum of A+B cups per wasp replicate was calculated). A few F_1_ individuals from each replicate were kept in -80°C to test for the presence of AnvRV by PCR. Replicates that did not have any female offspring were excluded from the analysis.

As the number of F_1_ female offspring in the RV^-^ line was very low in the developmental time (‘A’) cups, an additional experiment was carried out to test for this parameter (developmental time exp. #2). This experiment was set similarly to the first one, except that instead of placing 20 replicates of one pair of wasps, four replicates of five pairs of wasps per replicate were set, each cup provided with *ad-libitum* mealybugs.

Statistical analysis for all experiments was done using R software by a student t-test, except for the longevity experiment which was analyzed by a Kaplan-Meier test.

### Superparasitism and encapsulation experiment

As mentioned in the introduction, encapsulation of the invading egg by the host and superparasitism behavior may both be modulated by viruses. This experiment was conducted to test whether AnvRV induces or affects both phenomena. Measuring encapsulation rates directly proved to be challenging, therefore it was evaluated using two indirect measures: i) the number of melanized capsule marks on eggs oviposited into mealybugs, *i*.*e*. black spots of melanin deposition formed around the wasp egg. This is not a direct measure, as melanin spots can also be triggered by other factors, such as ovipositor probing through the cuticle without egg deposition; ii) hatching rates of wasp eggs, *i*.*e*., proportion of eggs that presumably overcame possible encapsulation by the mealybug host. Individual *A. vladimiri* females (*n*=20 of both RV^-^ and RV^+^ lines) were let to parasitize 15 *P. citri* adult females for 48h. Then, the wasps were removed, the mealybugs were incubated for three additional days, then gently washed with a fine brush dipped in soap water to remove waxy material, and the number of melanized capsule marks that were visible through the washed cuticle were counted under a stereomicroscope (encapsulation measure #1) (Fig.2, A-C). Next, the number of eggs laid in each mealybug (superparasitism measure) and the presence of developing wasp larvae were recorded (*i*.*e*., hatching rates; proportion of eggs that overcame possible encapsulation). For that, the mealybugs were submerged in 90% lactic acid and heated to 80°C for 90min ^23^. This treatment makes eggs and developing larvae visible under a light microscope. Slides were prepared with Hoyer’s medium and observed under a light microscope using up to x200 magnification (Fig. 2, D-F).

**Fig. 2.**
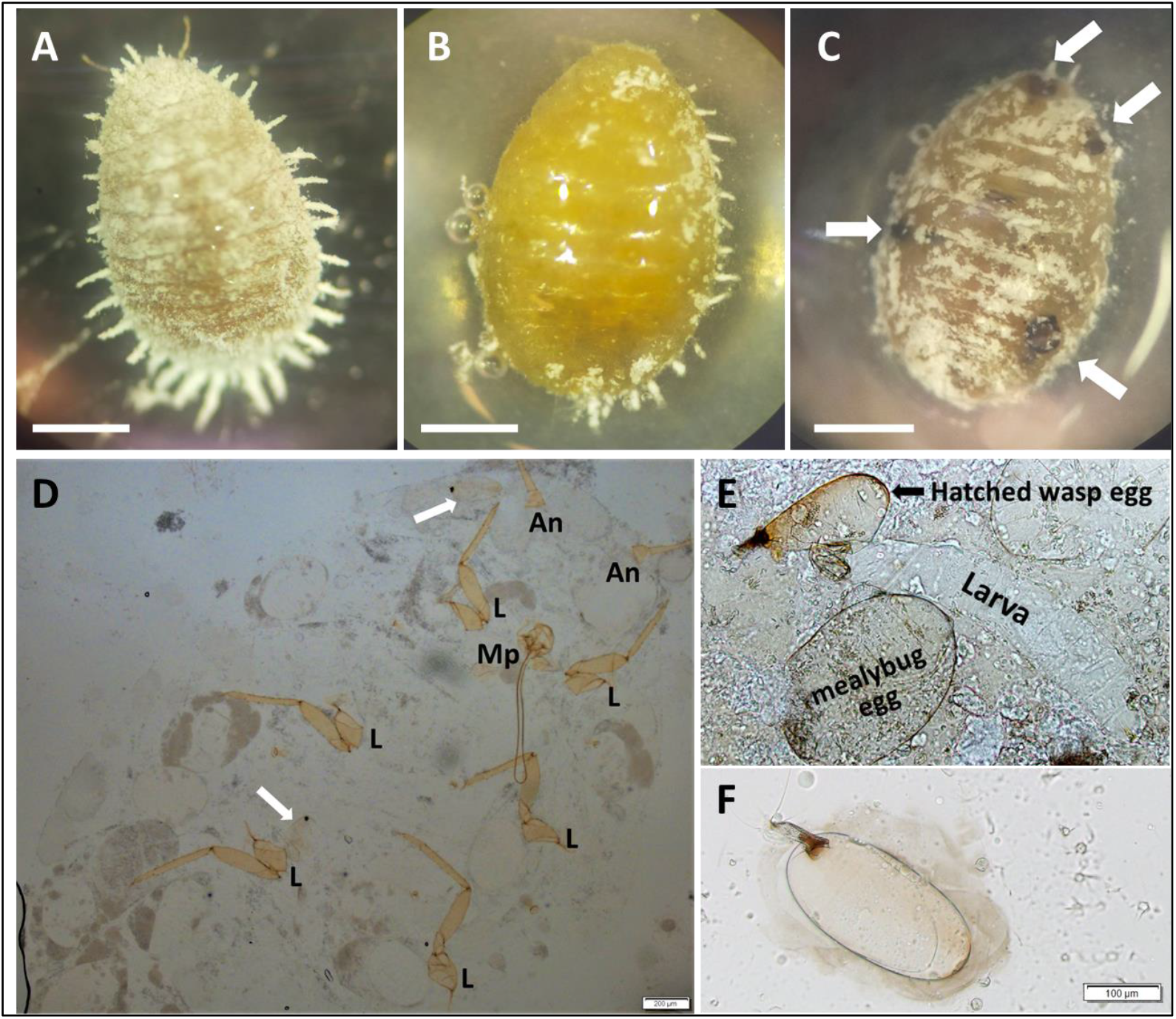
Capsule marks, wasp eggs and wasp larvae in parasitized mealybugs. A-C) Mealybugs before dissection, observed under a stereomicroscope, dorsal view. Scale bars, 1 mm. A) no treatment. B) washed with soap water. C) two days post parasitization, washed with soap water. White arrows point at capsule marks that can be seen through the washed cuticle. D-F) Dissected parasitized mealybugs cleared in lactic acid, observed under a light microscope. D) ventral view, whole body. White arrows point at wasp eggs. An, antenna. L, mealybug leg. Mp, mouth parts. Scale bar, 200 µm. Magnification X40. E) hatched wasp egg, and developing wasp larva inside a mealybug. Magnification X200. F) unhatched wasp egg. The surrounding dark ‘cloud’ may indicate the beginning of an encapsulation process. Scale bar, 100 µm. Magnification X200.

To assess superparasitism, the following measures were obtained: total number of eggs laid by each wasp in the 15 hosts, number and percent of parasitized mealybugs, and the average number of eggs laid per parasitized mealybug. To assess hatching rate values the calculation was: ((number of hosts containing hatched larvae/number of total hosts in the box (usually 15, unless some died)) × 100% (Pang et al., 2023).

To test whether AnvRV affects the hatching rate of *A. vladimiri* eggs (encapsulation measure #2), the number of parasitized mealybugs in which at least one wasp egg hatched was counted (*i*.*e*., at least one larva was found in the dissected mealybug). The hatching rate is portrayed in three ways, according to the number of wasp eggs observed in mealybugs: one egg per host (‘no superparasitism’), two or more eggs per host (‘superparasitized’), and all parasitized hosts pooled (‘all’).

To further validate the results of the hatching rate, the experiment was repeated a few months later, with the same wasp lines, using a larger sample size (*n*=31 RV^-^ line and *n*=29 for the RV^+^ line). Also, the parameter of capsule marks was repeated by a similar experiment but this time each female wasp (*n*=15) was given only two mealybugs to oviposit in, therefore increasing the incidence of superparasitism due to low host availability (‘high oviposition pressure’).

Statistical analyses to compare the means of all measures were done by students t-tests, except the measures of hatching rate which were analyzed with a GLM binomial test (R-version 4.1.2).

## Results

### Vertical transmission of AnvRV

In examining the transmission of AnvRV from parents to offspring, we discovered near-complete <u>maternal</u> transmission but no <u>paternal</u> transmission of the virus. Forty-six out of 47 offspring of 14 Lab-RV^+^ mothers (crossed with either Lab-RV^+^ or Field-RV^-^ fathers) were AnvRV-positive. Conversely, AnvRV could not be detected in any of the 31 offspring of eight Field-RV^-^ mothers crossed with Lab-RV^+^ fathers (Table 1).

**Table 1.** Vertical transmission of AnvRV in *Anagyrus vladimiri*. F=females, M=males.

| **Cross** | **F<sup>+</sup>M<sup>+</sup>** | **F<sup>+</sup>M<sup>-</sup>** | **F<sup>-</sup>M<sup>+</sup>** | **F<sup>-</sup>M<sup>-</sup>**<sup>481</sup> |
| --- | --- | --- | --- | --- |
| **Number of parent pairs** | 8 | 6 | 8 | 5 |
| **F1 tested** | 36 | 11 | 31 | 23 |
| **F1 positive** | 35 | 11 | 0 | 0 |
| **Transmission rate** | 97% | 100% | 0% | 0% |

### Horizontal transmission (HT) of AnvRV and establishment of RV^+^and RV^-^lines with identical genetic background

No HT occurred between <u>adult</u> *A. vladimiri* in our experiment, as the number of infected wasps at the beginning and at the end of the experiment was identical (see methods). Conversely, horizontal transmission did occur between <u>larvae</u> developing together in a shared host, with an average probability of ∼23% (15 out of 64, summing results of the two experiments). Subsequently, the newly infected *A. vladimiri* females had transmitted the virus vertically to 22 out of 24 offspring (92%). Offspring of the F_2_ AnvRV-positive females were used as founders of a wasp line which has identical genetic background as that of the field collected RV^-^ line, termed hereafter ‘Field-RV_**+**_’ (or ‘RV^+^’) line (see Fig. 1).

### Effects of AnvRV on the fitness of A. vladimiri

There were no statistically significant differences between the RV^-^ and the RV^+^ lines in all basic fitness parameters tested, including developmental time (two experiments), longevity, fecundity (48h and lifetime), and sex ratio (Table 2).

**Table 2.**
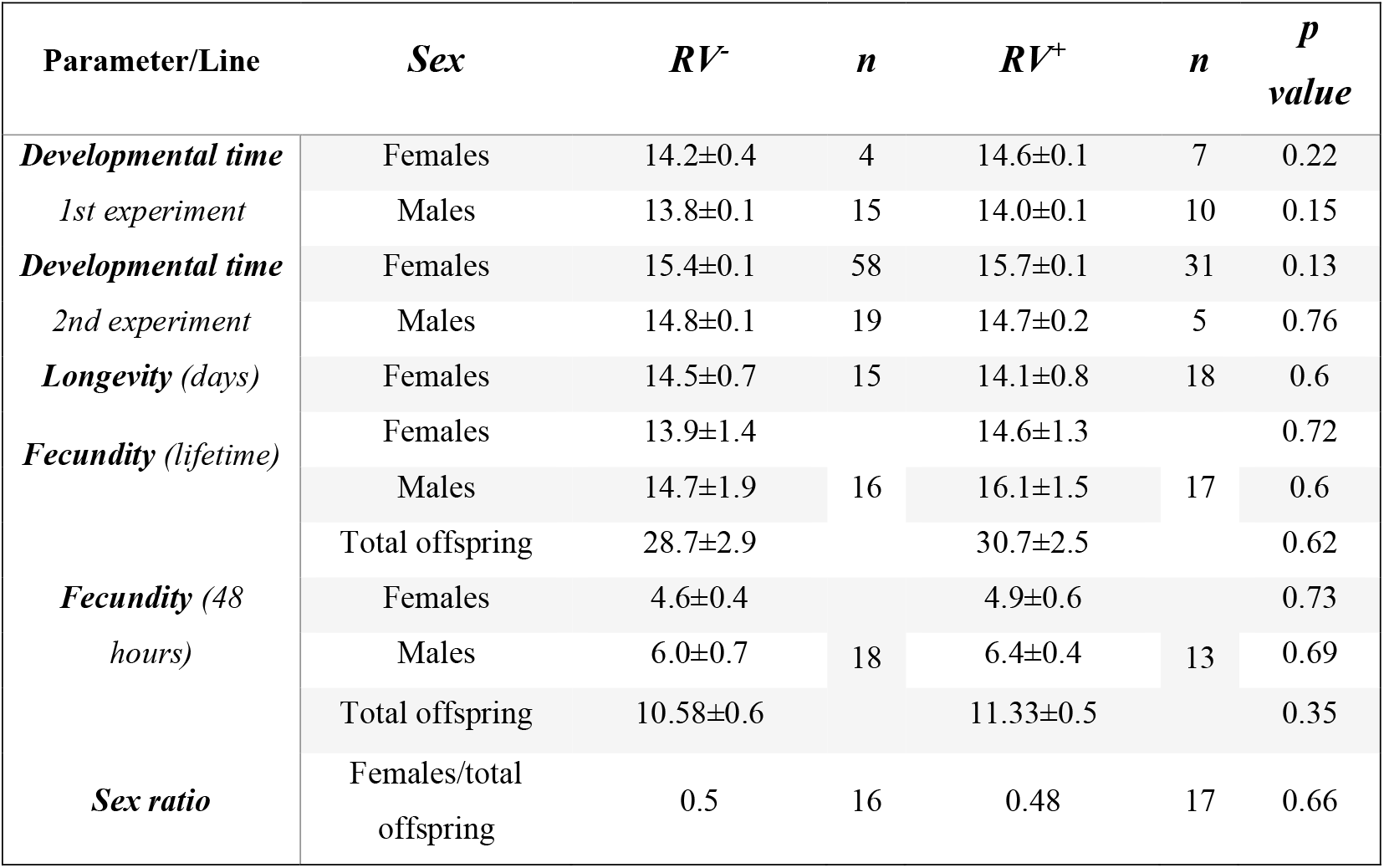
Various fitness parameters of *Anagyrus vladimiri* RV^-^ and RV^+^ lines. No significant differences were observed between the lines. P values are given for *t tests*, except for the longevity which was analyzed with a *Kaplan-Meier test*. Values are given as mean±standard error.

### Effect of AnvRV on superparasitism and encapsulation

In both experiments, there were no statistically significant differences between the wasp lines in the parameters of superparasitism, i.e., the number of wasp eggs laid per parasitized mealybug host (Table S1).

Conversely, the egg hatching rate and number of capsule marks differed significantly between the two strains. Although wasps from both lines laid on average the same number of eggs per host, significantly more wasp larvae were observed in mealybugs parasitized by the RV^+^ line compared to mealybugs parasitized by the RV^-^ line, *i*.*e*., the hatching rate of RV^+^ *A. vladimiri* is significantly higher (Fig. 3). The difference in hatching rate was most conspicuous when only singly parasitized hosts were compared (*i*.*e*., only mealybugs with a single *A. vladimiri* egg; 84.3% vs. 37.8%, Binomial GLM, p<0.0001; Fig. 3A). When only superparasitized mealybugs were compared, egg hatching rates were high for both lines, but still 9.5% higher in the RV^+^ line (Binomial GLM, p=0.035; Fig. 3B). When all parasitized mealybugs were compared, the hatching in the RV^+^ wasps was 18% higher than in the RV^-^(Binomial GLM, p=0.00025; Fig. 3C).

**Fig. 3.**
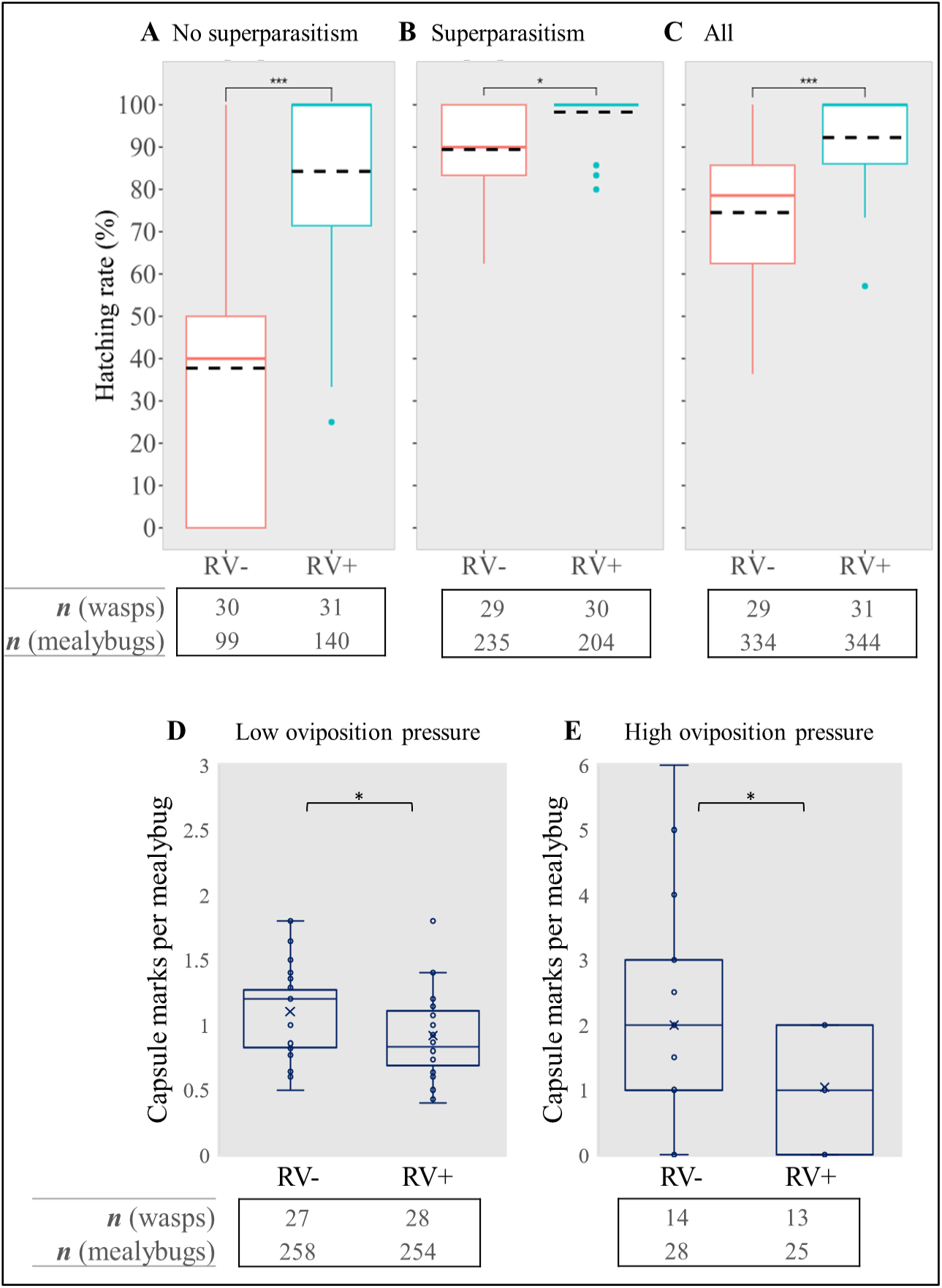
Egg hatching rate and encapsulation of wasp eggs by the mealybug host. **A-C**; Egg hatching rate of RV^-^ and RV^+^ *A. vladimiri* **- A**) no superparasitism: only mealybugs with one *A. vladimiri* egg or larva are counted; **B**) with superparasitism: only mealybugs with more than one *A. vladimiri* egg or larva(e) are counted; **C**) all parasitized mealybugs counted. Black dashed lines indicate the averages. **D-E)** number of capsule marks formed by mealybugs parasitized by *A. vladimiri*; **D)** in ‘low oviposition pressure’, each wasp had 15 mealybugs to oviposit in for 48 hours, and **E)** in ‘high oviposition pressure’, each wasp had only two mealybugs to oviposit in for 48 hours. Blue ‘x’: average, straight line: median.

Similar significant differences were also obtained when the experiment was repeated independently with the same conditions (but smaller no. of replicates). The hatching rates of the RV^+^ eggs were higher than those of the RV^-^ eggs by ∼30% in the total replicates of mealybugs (Binomial GLM, p=0.0015), by ∼40% in ‘no superparasitism’ (Binomial GLM, p=0.00036), and by ∼25% ‘with superparasitism’ (Binomial GLM, p=0.026) (Fig. S1).

The number of capsule marks also differed significantly between the lines: mealybugs parasitized by the RV^-^ wasps had 1.1±0.06 encapsulation marks, and those parasitized by RV^+^ wasps had 0.9±0.06 (t_56_=2, p=0.021; Fig. 3D). An even larger difference in this parameter was found when only two mealybugs were given per wasp (higher oviposition pressure): 2.0±0.3 encapsulation marks in the RV^-^ line, vs. 1.04±0.2 in the RV^+^ line (t_26_=2.06, p=0.022; Fig. 3E).

In summary, AnvRV affects both hatching rate and capsule marks. In both cases this effect was consistently detected in two independent experiments.

## Discussion

The current study presents a unique example of an RNA virus that increases the probability that the egg of its parasitoid wasp host will hatch. This finding was made possible only after determining the virus’s modes of transmission and harnessing the horizontal mode of transfer to introduce the virus into previously uninfected wasps, effectively establishing new wasp lines that could be used in phenotypic bioassays.

Our experiments show the major effect of AnvRV on wasps’ egg hatching rates, particularly when mealybugs were parasitized only once (Fig. 3A). In superparasitized mealybugs the differences between the wasp lines were less dramatic because the egg hatching rates of RV^-^ wasps increased significantly (Fig. 3B). This is expected: the more eggs laid in the host, the more likely that at least one of them will not be encapsulated (Suma et al., 2012), therefore superparasitism may be an adaptive strategy to compensate for eggs eliminated by the host and improve parasitism success in this species. Similar results were obtained on a related system (Sagarra et al., 2000), suggesting this saturating effect of superparasitism is at play in pseudococcidae/Encyrtidae interactions in general and possibly in other host-parasitoid systems as well.

One would expect that the positive effect of AnvRV on egg hatching will translate to higher fecundity of RV^**+**^ wasps than RV^**-**^wasps. However, the total offspring numbers were found to be very similar in the two lines. Because encapsulation may be exerted on the parasitoid larvae rather than the eggs (Blumberg, 1997), one possibility is that AnvRV’s counter-encapsulation effect observed on eggs is cancelled out by an increase of larvae encapsulation. Additionally, sometimes eggs survive encapsulation even though they seem to be fully melanized ^19^, therefore, perhaps the capsule marks measure does not fully indicate successful elimination of the parasitoid.

Variations in encapsulation levels and survival rates have been noticed among different ecotypes of *A. vladimiri*, for example, ∼15% encapsulation in an ecotype from Israel (Blumberg et al., 1995) vs. ∼60% encapsulation in an ecotype from Sicily (Suma et al., 2012). Knowing the interplay between AnvRV presence and the mealybug’s immune system, it would be interesting to test whether these substantial differences in encapsulation rates are correlated with the prevalence of AnvRV in these populations.

### Transmission routes of AnvRV

Our study is the first to show a Reovirus using both VT and HT. Generally, Reoviruses in insects are transmitted either transovarially (Juchault et al., 1991; López Ferber et al., 1997; Meng et al., 2019) or horizontally through the ingestion of infected feces (Renault et al., 2005; Matthijnssens et al., 2022). The mixed transmission mode of AnvRV is efficient: high-fidelity VT maintains the virus within a lineage, while HT allows it to spread under conditions of high parasitoid density (Bailly-Bechet et al., 2017). Horizontal transmission (HT) of viruses plays a key role in virus spreading, particularly under conditions of high parasitoid / host ratios (e.g. (Cory, 2015). In environments with high competition for hosts superparasitism increases, facilitating HT. For example, AnvRV is fixed in the *A. vladimiri* lab strain used here and in our previous study (Izraeli et al., 2022), but far less prevalent in field populations (three RV^+^ wasps out of 24), possibly because reduced competition limits HT opportunities. Hence, AnvRV may spread in a population by HT and subsequently persist in infected lineages by maternal transmission. Studies on other parasitoids, like *Leptopilina boulardi* (Patot et al., 2010), have shown similar patterns, where denser populations promote viral HT.

A virus benefiting from both horizontal and vertical transmission may increase in frequency because of sufficient opportunities for HT, and/or because it increases its host fitness (i.e. through VT). We found that AnvRV was associated with a higher hatching rate of larvae, opening the possibility that AnvRV increases wasp fitness. If this is indeed the case, AnvRV should go to fixation in natural populations. On the contrary, viral prevalence was low in natural populations of *A. vladimiri*, thus questioning this interpretation. This apparent discrepancy may come from the fact that our experiments were conducted on *P. citri*, while the main host in the natural population was *P. ficus*. Thus, we can still speculate that AnvRV increases *A. vladimiri* fitness only in specific contexts, such as when parasitizing specific mealybug host species, or by conferring tolerance to abiotic factors, or protecting from hyperparasitoids (Vorburger, 2022). If the fitness boost only applies to less common hosts in the field, this could explain the moderate viral prevalence observed in natural populations. Host-symbiont interactions in other insects (Sanaei et al., 2021; Kumar Pradhan et al., 2024) support this explanation, and further research on additional mealybug host species could clarify AnvRV’s ecological role.

Finally, utilizing HT to introduce the virus into virus-free females through superparasitized mealybug hosts proved to be an effective method for establishing a virus-infected wasp line with the same genetic background as the virus-free line. Unlike microbial symbionts, which can be relatively easily “cured” from wasps using antibiotics, eliminating viruses is far more challenging. The success of this method should therefore be considered for the establishment of isogenic wasp lines when studying other virus-parasitoid systems.

### Consequences to biological control and conclusions

Despite being proposed several decades ago (Greany et al., 1984) ^41^ and emphasized in following publications (Zindel et al., 2011; Cusumano and Volkoff, 2021; Morrow et al., 2023), information on inherited viruses is still rarely applied to biocontrol. Instead, most studies have focused on plant, fungal, and insect pathogens (e.g. (Cusumano and Volkoff, 2021; Gutiérrez-Cárdenas et al., 2023).

The current study presents a unique natural enemy-virus system that has high potential for application in improvement of biocontrol of a devastating agricultural pest. The possible contribution of AnvRV to the fitness of *A. vladimiri* should be further investigated under field conditions, and with other host species, such as the vine mealybug, *P. ficus*. Also, the abundance of potentially beneficial viruses in taxonomically related parasitoid species should be determined. If similar beneficial effects are found, such information may be applied for improving the biocontrol of mealybug pests.

## Acknowledgments

The authors would like to thank Mr. Eyal Erel from ‘BioBee Sde Eliyahu Ltd’ for insect consultancy, Ms. Shira Gal from the ARO for visualization assistance with light microscopy and Mrs. Gefen Zomer for laboratory help. This research was supported by the Israel Science Foundation (grant No. 397/21) to E.Z-F. and E.C.

