## supplement information for "A Vertically and Horizontally Transmitted RNA Virus Facilitates Egg Hatching of a Parasitoid Wasp"

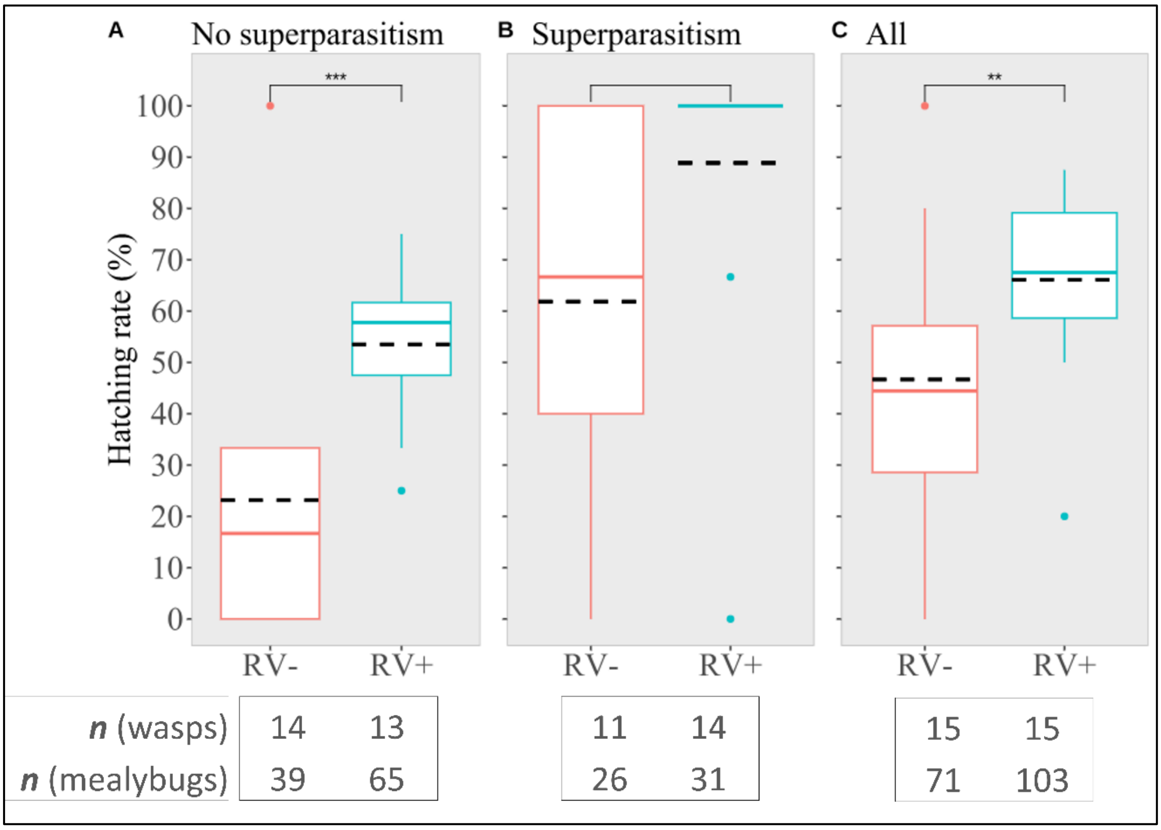


**Fig. S1. Egg hatching rate of RV$^-$ and RV$^+$ *Anagyrus vladimiri*. 1$^{st}$ experiment (smaller number of replicates). A**) no superparasitism: only mealybugs with one *A. vladimiri* egg or larva are counted; **B**) with superparasitism: only mealybugs with more than one *A. vladimiri* egg or larva(e) are counted; **C**) total replicates of parasitized mealybugs counted.

**Table S1. Superparasitism behavior compared between the $RV^-$ and $RV^+$ lines of *Anagyrus vladimiri*, which were allowed to oviposit in 15 mealybugs for 48 hours.** Results obtained in two independent experiments. Values are given as mean±standard error.

| Experiment 1 | | | |
|---|---|---|---|
| ***Parameter/Line*** | **$RV^+$** | **$RV^-$** | **P-value** |
| *n* | 19 | 15 | |
| *Eggs per wasp* | 7.42±1.1 | 7.4±1.3 | 0.99 |
| *Number of parasitized mealybugs* | 5.58±0.8 | 4.73±0.6 | 0.4 |
| *Percent of parasitized mealybugs* | 43.22±5.3 | 35.09±4.2 | 0.24 |
| *Number of eggs per parasitized host* | 1.32±0.06 | 1.44±0.1 | 0.31 |
| Experiment 2 | | | |
| ***Parameter/Line*** | **$RV^+$** | **$RV^-$** | **P-value** |
| *n* | 31 | 29 | |
| *Eggs per wasp* | 20.6±1.19 | 23.1±1.2 | 0.14 |
| *Number of parasitized mealybugs* | 11.1±2.5 | 11.5±2.0 | 0.48 |
| *Percent of parasitized mealybugs* | 79.9±2.8 | 83.1±2.0 | 0.90 |
| *Number of eggs per parasitized host* | 1.8±0.36 | 2.0±0.36 | 0.08 |
